# SCV-2000bp: a primer panel for SARS-CoV-2 full-genome sequencing

**DOI:** 10.1101/2020.08.04.234880

**Authors:** AS Speranskaya, VV Kaptelova, AV Valdokhina, VP Bulanenko, AE Samoilov, EV Korneenko, OY Shipulina, VG Akimkin

## Abstract

Here we provide technical data for amplifying the complete genome of SARS-CoV-2 from clinical samples using only seventeen pairs of primers. We demonstrate that the СV2000bp primer panel successfully produces genomes when used with the residual total RNA extracts from positive clinical samples following diagnostic RT-PCRs (with Ct in the range from 13 to 20). The library preparation method reported here includes genome amplification of ~1750-2000 bp fragments followed by ultrasonic fragmentation combined with the introduction of Illumina compatible adapters. Using the SCV2000bp panel, 25 complete SARS-CoV-2 virus genome sequences were sequenced from clinical samples of COVID-19 patients from Moscow obtained in late March - early April.

## INTRODUCTION

Presently, the SARS-CoV-2 pandemic has been raging in every country in the world, since it started in China in late 2019. The primary objective of epidemiological surveillance during the pandemic is to track its geographical spread, and this activity should continue throughout the pandemic. Analyzing the genome of SARS-CoV-2 from clinical samples is crucial for the understanding of viral spread and evolution as well as for timely developing epidemiology response strategies. Achieving these goals locally and nationally requires genome sequencing methods development.

NGS is the current mainstream method for SARS-CoV-2 genome sequencing. One of the most widely used approaches is the ARTIC protocol which includes 98 pairs of primers divided into two multiplex PCR reactions and covers almost the whole region of SARS-CoV-2 genome [1,2]. ARTIC protocol is popular, and its existing modifications improve the coverage of sequencing regions [3]. Nevertheless, it has some irreparable drawbacks, for example, a lot of short amplification fragments; as a result, it has a reduced ability to amplify the degenerative variants of virus genomes. [4]. Another drawback is a large number of oligonucleotides (totally 196 primers), so the total price for their synthesis could significantly limit the capabilities of small local laboratories for the production of genomic data due to the high input cost of experiments. There is a need for a decrease of the cost of library preparation for high-throughput sequencing.

Here we present a new primer panel which allows amplifying the complete genome of SARS-CoV-2 (the causative virus of COVID-19) using 17 primer pairs (in four pools). Our results demonstrate that our method allows producing full genomes when we use RNA extract from SARS-CoV-2 positive clinical samples which have a cycle threshold (Ct) in the range of 13 to 26. The resulting primer set exhibits the coverage of the entire viral genome except only 8 bp on 5’- and 80 bp on 3’- ends in comparison with the reference genome in GenBank (accession number MT121215.1).

## MATERIALS AND METHODS

### Clinical samples and RNA extraction

The swab samples from patients with symptoms of acute respiratory infections or suspected for SARS-CoV-2 were processed at the Molecular diagnostic laboratory of Federal Budget Institution of Science “Central Research Institute of Epidemiology” of The Federal Service on Customers’ Rights Protection and Human Well-being Surveillance (the laboratory licensed for working with pathogens) using the AmpliSens® Cov-Bat-FL assay kit based on real-time RT-PCR, detecting SARS-Cov-Family, including new SARS-Cov-2 (CRIE) test according to the manufacturer’s instructions. For genomes amplification, we used the remains of RNA samples extracted from oropharyngeal swabs in the clinical lab. The only chosen samples were those where the presence of SARS-CoV-2 was determined with a cycle threshold (Ct) <= 20. Before the work started, these RNA samples had been stored carelessly in the clinical lab under non-specialized conditions for 2-3 months.

### Primer design for SARS-CoV-2 complete genome multiplex PCR

The primer scheme for amplifying the whole genome of SARS-CoV-2 was initially designed using a web-based primer design tool – Primal Scheme [5, 6]. As a result, 17 primer pairs were designed. Thirty most variable SARS-CoV-2 full genome sequences available at the GenBank as of 30.03.2000 were used as reference genomes. The parameter of amplicon lengths of 2000 nts with a 100-nt overlap was used for the scheme generation. Then the designed primers were manually corrected and combined in pools. Subsequently, the primer combinations were optimized using iterative sequencing of some randomly selected samples on Illumina MiSeq. Finally, the designed primers were mixed equally into four pools, each containing from 2 to 6 primer pairs (see Table 1).

**Table 1.**
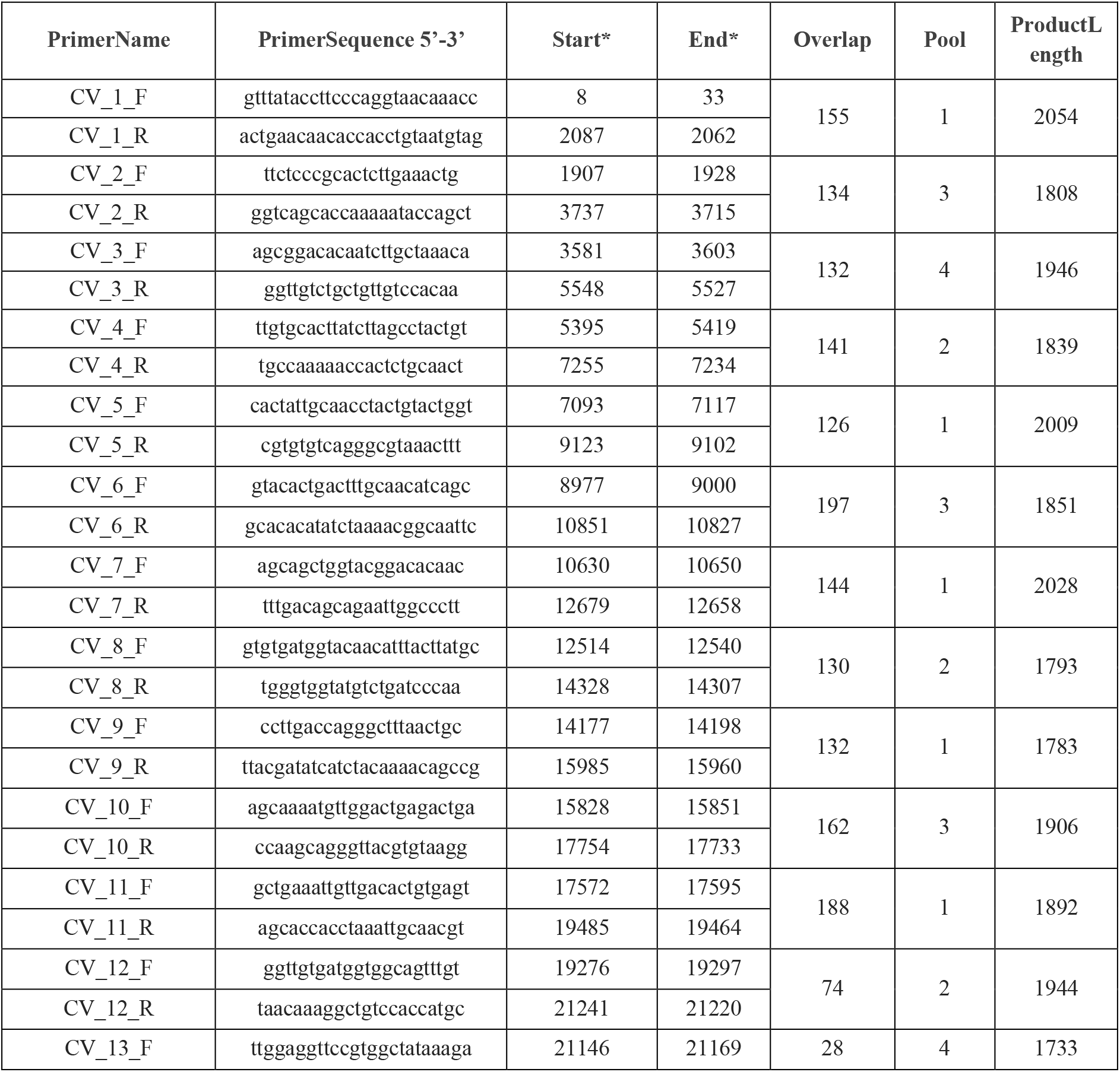
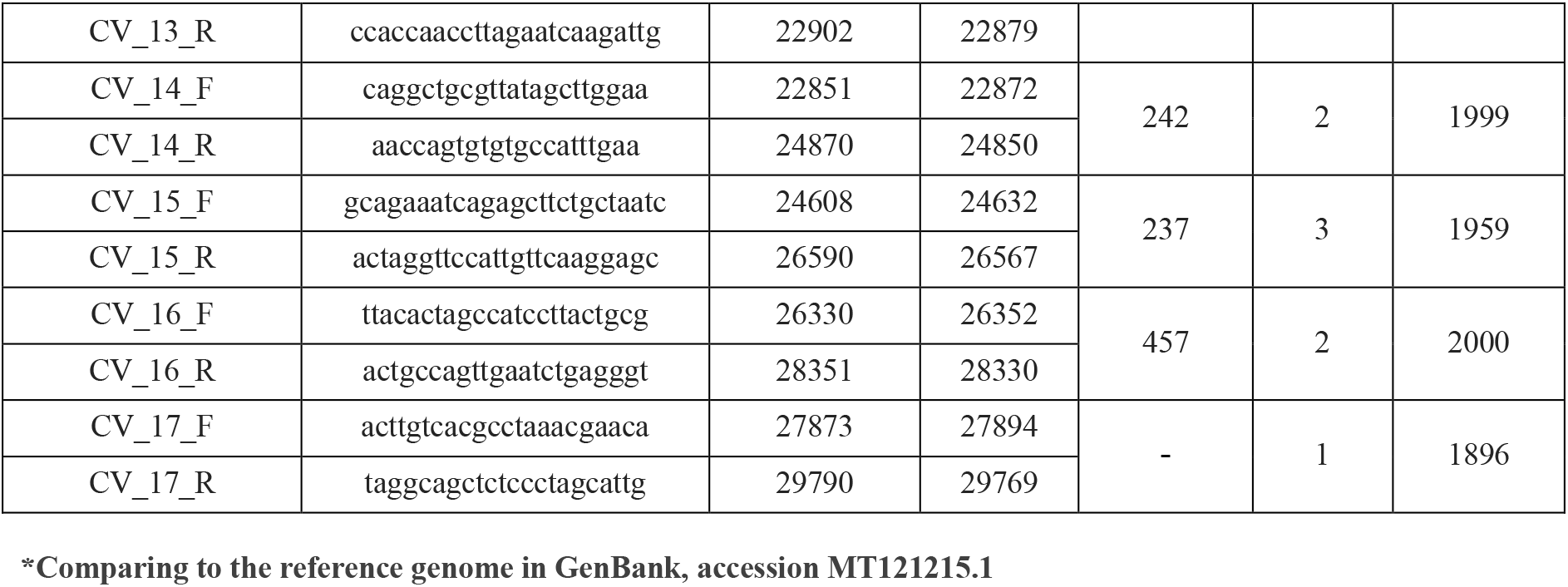
Primers for whole-genome sequencing of the SARS-CoV-2.

### Library preparation

Reverse transcription reaction was performed using 10 μL of the RNA samples, random hexanucleotide primers and Reverta-L kit (AmpliSens, Russia) according to the manufacturer’s instructions. The cDNA was immediately used as a template for the amplification of genome fragments.

The ~2000 bp SARS-CoV-2 genome fragments were amplified using the designed primers. The multiplex PCR amplification reaction was performed in the volume of 25-μL, containing 5-8 μL of the template cDNA (see **Table 2.**), 2 μL of the pooled primers (for the concentration see **Table 2.**), 0,5 μL of dNTPs (10 mM, Thermo Fisher Scientific, USA), Q5 High-Fidelity DNA Polymerase and polymerase buffer according to the manufacturer’s instructions (New England BioLabs, NEB). The PCR amplifications were performed using 96 well plates and Veriti™ 96-Well Thermal Cycler (Thermo Fisher Scientific).

**Table 2.** Concentrations of pooled primers

| Pool name | Quantity of | Total concentration | Concentration of each | Volume of cDNA |
| --- | --- | --- | --- | --- |

| | primers | ,$\mu\text{M}/\mu\text{L}$ | primer in pool per<br>PCR reaction, $\mu\text{M}/\mu\text{L}$ | per PCR reaction,<br>$\mu\text{L}$ |
| --- | --- | --- | --- | --- |
| 1 | 12 | 150 | 25 | 5 |
| 2 | 10 | 100 | 20 | 8 |
| 3 | 8 | 50 | 12,5 | 5 |
| 4 | 4 | 50 | 25 | 5 |

PCR conditions were the following: initial denaturation at 98°C for 30 sec, followed by 35 cycles of 98°C for 10 sec, annealing temperature at 64°C for 30 sec, 72°C for 2 min and final extension at 72°C for 2 min.

We used the following approach: at first we amplified only pool 2 because this pool works worst of all. Then products of the amplification reaction were analyzed with electrophoresis using 1,7% agarose gel stained with SYBR Green. The samples in which visible PCR products of the expected size were found were used for PCR reactions with other primer pools (1, 3 and 4). Then products of PCR were visualized using 1,7% agarose electrophoresis stained with SYBR Green and amplified fragments were mixed in equimolar amounts according to visual estimation of concentration. To purify PCR products of the expected size (1700-2100 bp) from the reaction mixture and to remove the nonspecific short fragments obtained during the amplification step, premixed amplicons were cleaned in the ratio 0,7x using carboxyl-coated magnetic particles, commercially available as Agencourt AMPure XP purification system (Beckman Coulter, Danvers, MA, USA). The concentrations of the purified fragments were measured using Qubit ds DNA HS Assay Kit with a Qubit 3.0 fluorometer (Invitrogen).

### Ligation-based library preparation

The premixed and purified PCR products (of 17-300 ng per sample) were sheared in microTUBE-50 AFA Fiber Screw-Cap (PN 520166) using Covaris M220 (Covaris, Woburn, MA) using the following settings: peak incident power (W)- 75, duty factor- 5%, cycles per burst-200, treatment Time (s) - 50, temperature (°C)-20, sample volume (μl)- 50. Paired-end libraries were constructed with NEBNext® Ultra™ End Repair/dA-Tailing Module (NEB E7442L), NEBNext® Ultra™ Ligation Module (NEB E7445L) and Y-shaped adapters compatible with Nextera XT Index Kit. The amplification of libraries was performed with Q5 High-Fidelity DNA Polymerase in 25 μL according to the manufacturer instructions (New England BioLabs, NEB) with 10 cycles of amplification.

### Tagmentation-based library preparation

Also, another method was used for library preparation, namely the workflow of Nextera™ Library Prep (Illumina, USA). For library preparation, 50 ng of mixed and purified PCR products were used. The libraries were created according to the Nextera protocol subject to the following changes: amplification was performed with Q5 High-Fidelity DNA Polymerase, dNTPs (10 mM, Thermo Fisher Scientific, USA) and intercalating dye EvaGreen (Biotium, USA) with 8 cycles of amplification.

The size selection of the final libraries was done using Agencourt AMPure XP (Beckman Coulter, Danvers, MA, USA). The quality and fragment length distribution of the obtained libraries were evaluated with Agilent Bioanalyzer 2100 (Agilent Technologies, USA). Sequencing was performed on Illumina HiSeq 1500 with HiSeq PE Rapid Cluster Kit v2 and HiSeq Rapid SBS Kit v2 (500 cycles) or Illumina MiSeq with the MiSeq Reagent Kit V2 (500 cycles).

### Data analysis

To process raw reads of multiple samples, we have utilized a pipeline consisting of mainstream bioinformatics tools. In short, reads were trimmed by quality with Trimmomatic [7]. PCR primers were removed using Cutadapt [8]. Resulting reads were mapped to reference genome MT121215.1 using bowtie2 [9]. Reads with low mapping quality were filtered out using SAMtools [10]. Basecalling was performed with GATK [11]. Gvcf files were filtered with BCFtools [12], and the consensus sequence was obtained with BEDTools [13]. Finally, the validity of the obtained sequence was verified manually by mapping reads to the consensus sequence followed by visual inspection. Areas with read coverage lower than 5 were masked with NNN.

## RESULTS

We have developed a multiplex system to amplify the long nucleic acid fragments of SARS-CoV-2 genome. The designed scheme of primers, named SCV2000bp, produces overlapping PCR products of 1733-2054 bp. The overlaps between different amplicons are 28-457 bp. The produced fragments cover the entire genome of SARS-CoV-2 with the exception of 7 bp at the 5’-end and 80 bp at the 3’-end comparing to the reference genome in GenBank, accession number MT121215.1 (without poly(A) tail).

The ligation adapters approach was used for the successful sequencing of 20 resulting samples. We got 1.57±0.76 million reads (631±288 mbp) of sequence data per sample. After removal of low-quality reads and primer trimming, we achieved the average coverage of the reference SARS-CoV-2 genome of 8800±4400. For the example of coverage see Figure 1. The complete genomes for SARS-CoV-2 were assembled with a custom script implementing the bioinformatics workflow, and these results were confirmed by CLC Genomic Workbench 8.5. Mutations in the regions with low coverage and primer-related regions were manually checked with the use of reference MT121215.1. The sequences were submitted in GISAID database with accession numbers: EPI_ISL_479620 - EPI_ISL_479624, EPI_ISL_486815 - EPI_ISL_486829.

**Figure 1.**
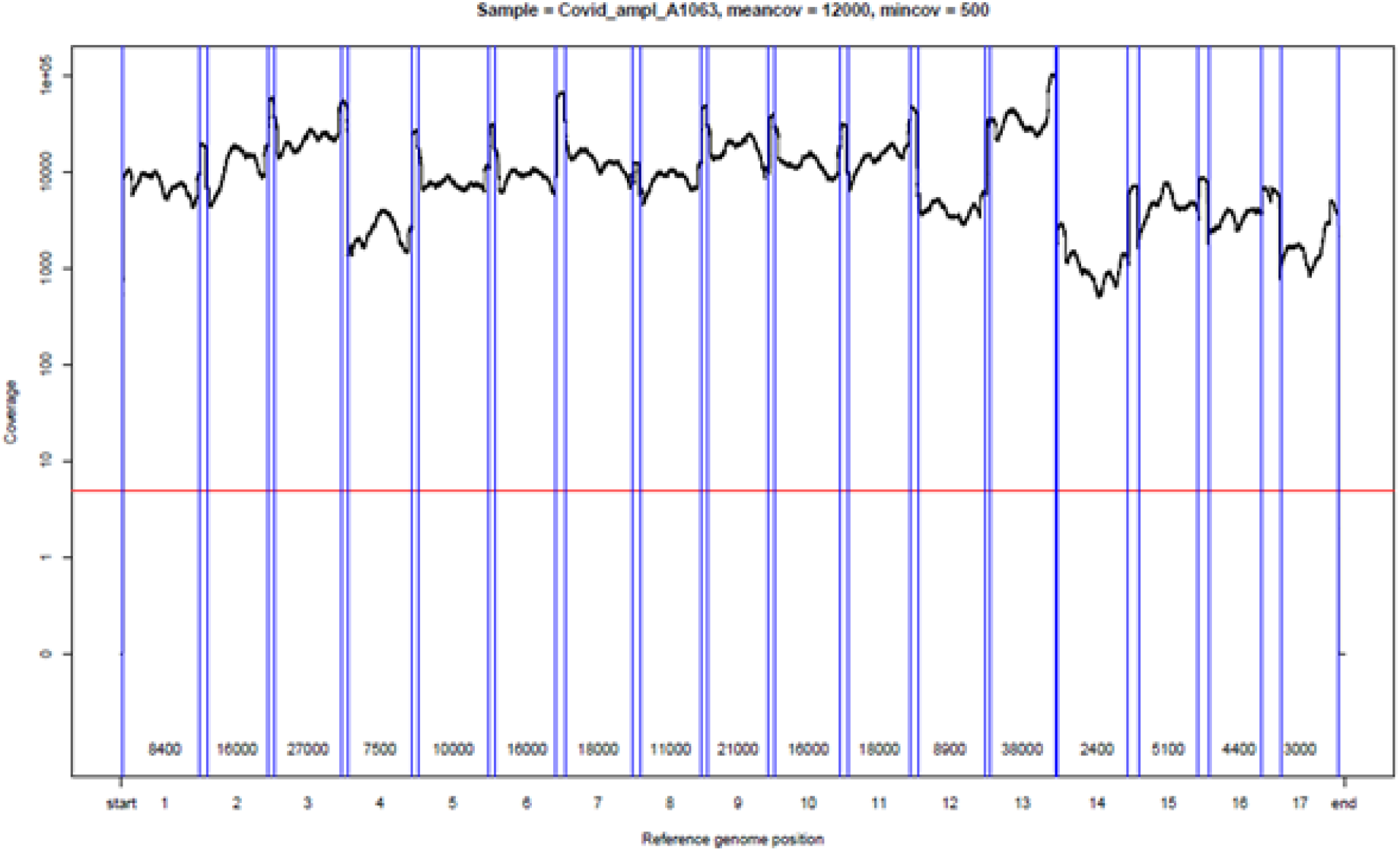
This picture demonstrates the results of the sequencing of sample A1063. The data were obtained using SCV2000bp primer panel and the ligation adapters approach. Here you can see the layout of the amplicons 1-17 along the reference genome (X-axis) and coverage (Y-axis).

The ligation adapters approach includes the stage of nucleic acids random fragmentation followed by adapter ligation. Unfortunately, this approach is rather expensive due to the cost of equipment for fragmentation and components of adapter ligation kits (ligase the most expensive). Therefore, we checked the feasibility of another tagmentation-based library construction method. For library preparation, we used SCV2000bp primer set and five SARS-CoV-2 positive samples. We sequenced libraries on MiSeq PE 250+250. The sequencing resulted in 303±84 thousand reads totalling 140±35 mbp of sequence data per sample. So, five whole SARS-CoV-2 genomes were successfully assembled. The average coverage of the genomes was equal to 3040±770. For the example of coverage see Figure 2. The sequences were submitted in GISAID database with accession numbers: EPI_ISL_467774, EPI_ISL_467775, EPI_ISL_470539, EPI_ISL_462149, EPI_ISL_462150.

**Figure 2.**
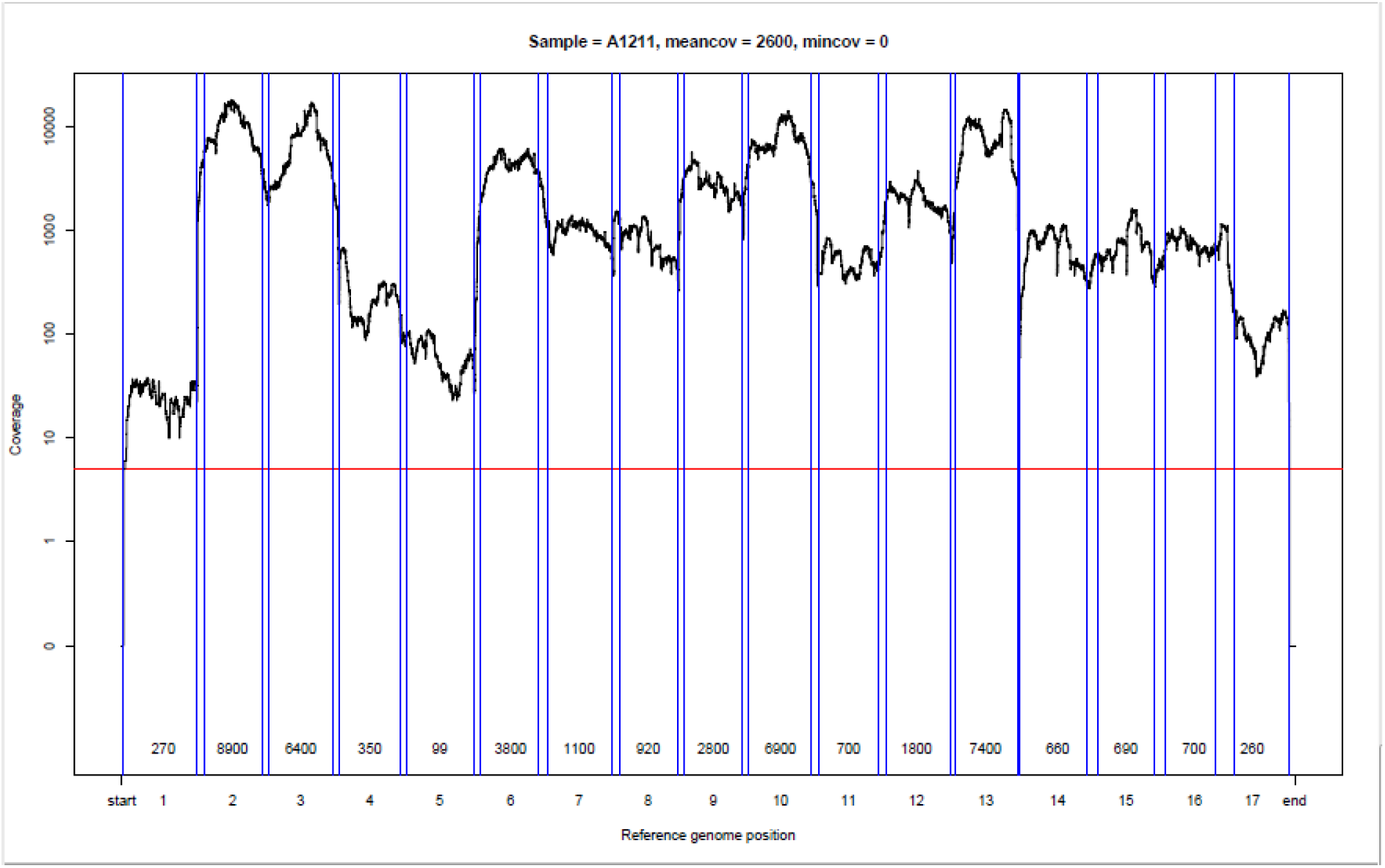
This picture demonstrates the results of the sequencing of sample 1211, using SCV2000bp primer panel and tagmentation-based approach. Here you can see the layout of the amplicons 1-17 along the reference genome (X-axis) and coverage (Y-axis).

The synthesizing of cDNA from input RNA using random primers followed by the polymerase chain reaction with specific primers is the most common method for amplifying RNA viruses to obtain sufficient coverage and depth of the whole viral genome. For NGS library preparation, we used the same RNA samples which were the remains of clinical diagnostic (i.e. without repeating the stage of RNA extraction). All the analyzed samples showed the Ct value in the range of 13 to 20 (i.e. a high virus load). Under the conditions described in Materials and Methods, multiplex PCR successfully produced fragments of the expected length. The amplification was generally less effective for pool 1 and pool 2 primers (most likely due to the presence of unspecific products) (See Figure 3).

**Figure 3.**
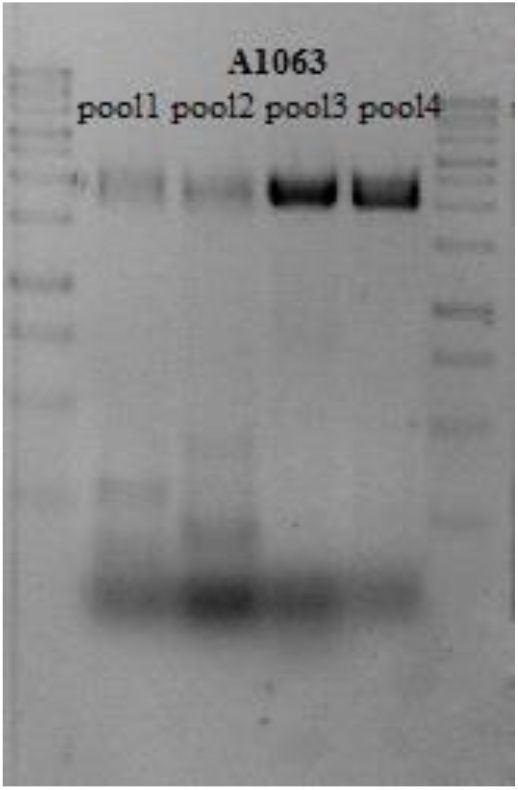
The primer pools testing using gel electrophoresis. The results of amplification of sample A1063 are demonstrated. A 1kb DNA ladder (NL001, Evrogen) used in the rightmost and leftmost lanes (250bp, 500bp, 750bp, 2×1000bp, 1500 bp, 2000bp, 2500bp, 2×3000bp, 4000bp, 5000bp, 6000bp, 8000bp, 10000bp).

The primer panel was tested with many different RNA samples, namely, we tested 233 samples with Ct from 13 to 26. Some samples failed in amplification with pool1, pool 2, and even with all four primer pools. Since our goal was to develop a method for sequencing viral genomes using the RNA samples remaining after diagnostic in the clinical labs, we tried to find a method for preliminary selection of promising samples directly from a great number of samples. Some authors claim that “poor quality consensus genomes were generally associated with a lower SARS-CoV-2 viral load in the clinical samples i.e. higher RT-qPCR Ct values” [14]. Therefore, we check the impact of Ct values differences on the successful genome amplification. We amplified only the second pool because it worked worst of all and checked the results of amplification using gel electrophoresis. We found that the number of successful amplifications is strongly related to the collection date and storage conditions of samples, but not to Ct value (See **Table 2**).

**Table 2.** SCV2000bp-amplification for the samples positive for SARS-CoV-2

| Ct | 13-17 | 17-20 | 21-26 | Alltypes |
| --- | --- | --- | --- | --- |
| **Lot A** |  |  |  |  |
| The number of samples | 20 | 15 | - | 35 |
| % of positive for pool 2 | 40% | 60% | - | 51% |
| % of positive for all four pools of primers | 35% | 46,7% | - | **46%** |
| **Lot B** |  |  |  |  |
| The number of samples | 26 | 50 | 58 | 134 |
| Of positive for pool 2 | 46% | 26% | 12% | 30,6% |
| of positive for all four pools of primers | 23% | 6% | 0% | **5,2%** |
| **Lot C** |  |  |  |  |
| The number of samples | 25 | 21 | 18 | 64 |
| of positive for pool 2 | 24% | 24% | 28% | 25% |
| of positive for all four pools of primers | 0% | 4,7% | 0% | **1,6%** |

Another reasonable explanation of the poor results of amplification was the degradation of the extracted RNA. So, we measured total RNA integrity number (RIN) using Agilent Bioanalyzer 2100 for a set of samples and found that successful amplification, sequencing and assembling could be achieved even for samples containing fully degraded RNA, see Table 3. The samples which are marked bold in Table 3, namely A1063, A1134, A1102, A1106 and A1076 were successfully amplified, sequenced and assembled, although their quality was extremely bad. And vice versa, even if RNA had RIN >5 (i.e. good enough quality), this did not guarantee the success of amplification. For example, see sample A1135, Table 3.

**Table 3.**
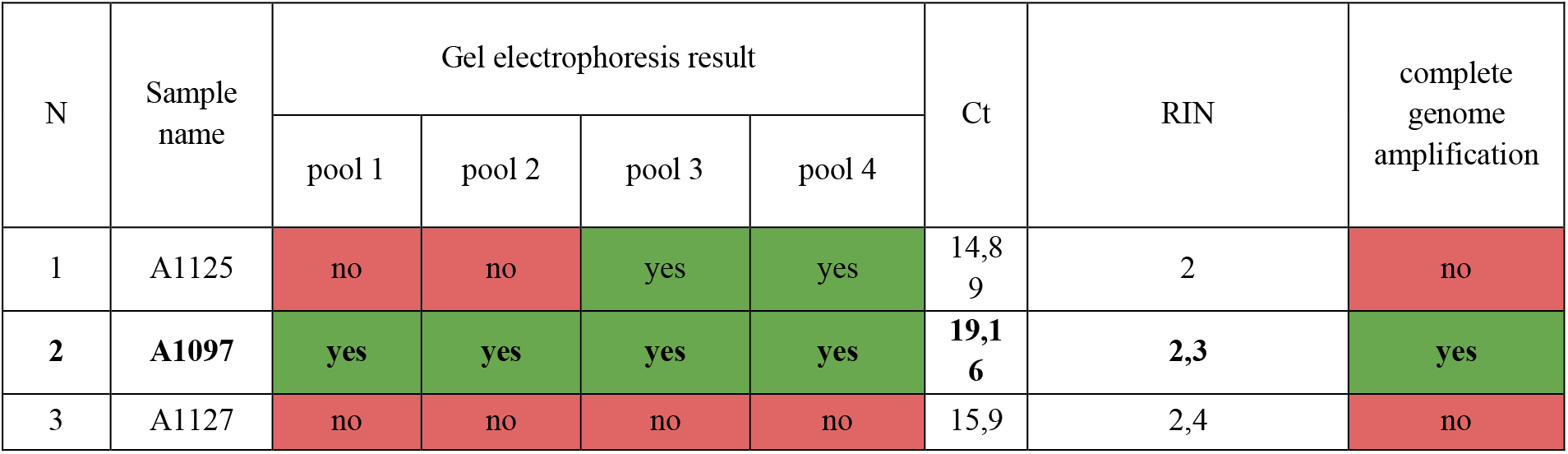
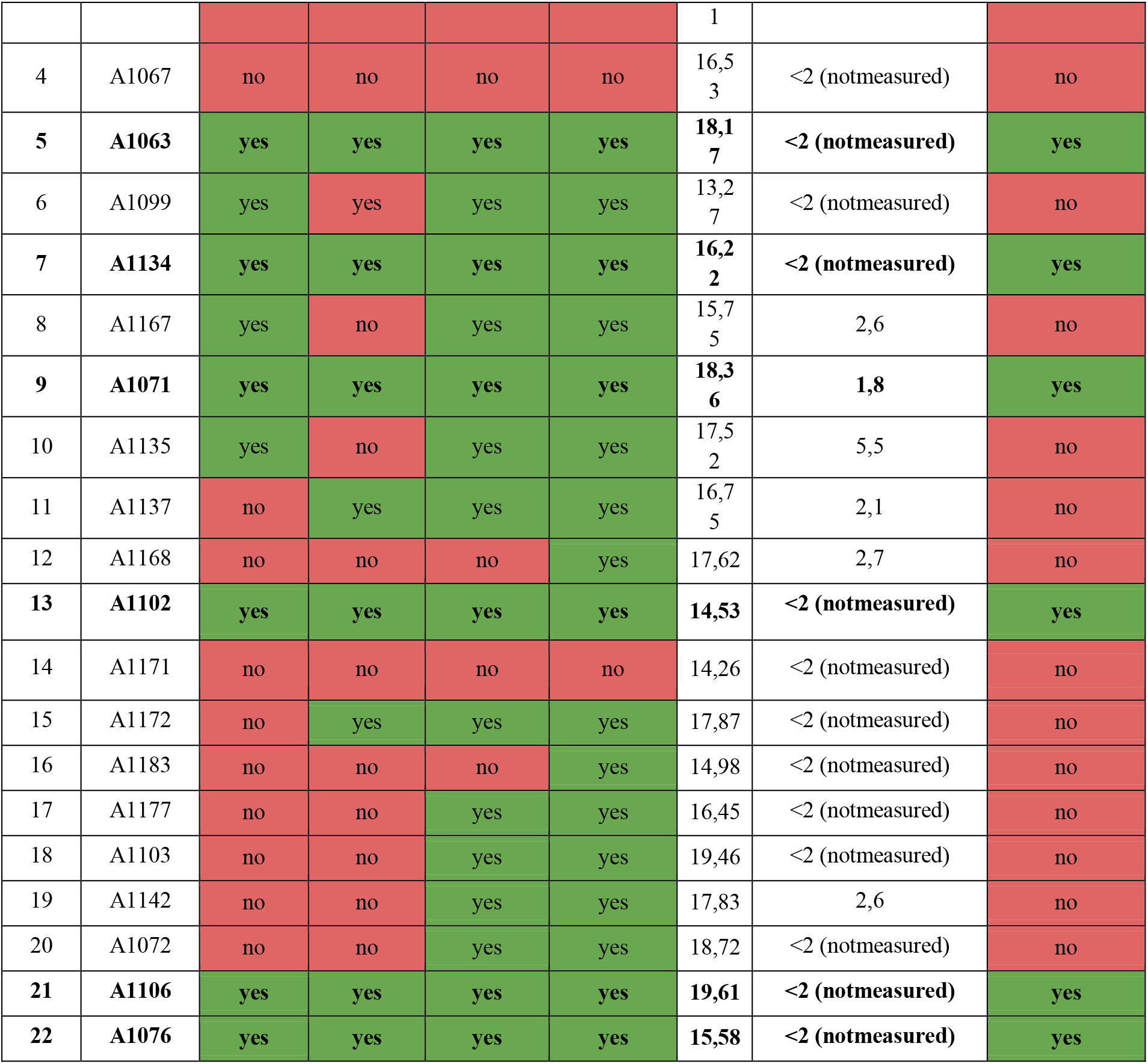
Total RNA degradation impact on the success of amplification, estimated by RIN.

## DISCUSSION/CONCLUSION

We developed a primer set for a relatively inexpensive generation of genomic data for SARS-CoV-2. Previously, a multiplexed PCR primer set for whole genome analysis of SARS-CoV-2 (ARTIC Network) was proposed by [1, 2, 3]. Currently, this is the most widely used approach. The cost of oligonucleotides synthesis for the ARTIC primer set is about 11980$ (if 5o.e. for each primer synthesized). So, the input price of experiments is high and could be a hindrance for researchers in small local labs with limited financial capabilities. Therefore, it is advisable to have a battery of methods for complete SARS-CoV-2 genome amplification. The cost of СV2000bp primer panel oligonucleotide synthesis is only 2300$. We demonstrate that this primer panel can be successfully used for sequencing and assembling of SARS-CoV-2 genomes both by ligation adapters approach and by tagmentation-based library construction if total RNA extracts from positive samples with Ct in the range from 13 to 20. Moreover, many research laboratories are not licensed to work with samples containing live SARS-CoV-2 viruses, so they have to deal with the remains of the RNAs extracted from oropharyngeal swabs in diagnostic labs. The library preparation methods described in this manuscript could be used for the complete genome sequencing of SARS-CoV-2 even from the samples which are remains of diagnostic tests in clinical labs.

